# A 4D Bioprinting Platform to Engineer Anisotropic Musculoskeletal Tissues by Spatially Patterning Microtissues into Temporally Adapting Support Baths

**DOI:** 10.1101/2025.05.01.651636

**Authors:** Francesca D. Spagnuolo, Gabriela S. Kronemberger, Daniel J. Kelly

## Abstract

Bioprinting is a powerful tool for engineering living tissues, however replicating native composition, structure and function remains a major challenge. During morphogenesis, cellular self-organization and matrix development are strongly influenced by the mechanical constraints provided by surrounding tissues, suggesting that such biophysical cues should be integrated into bioprinting strategies to engineer more biomimetic grafts. Here we introduce a novel 4D bioprinting platform that spatially patterns mesenchymal stem/stromal cell (MSC)-derived microtissues into temporally adapting support baths. By modulating the bath’s mechanical properties, we can precisely control the physical constraints applied post-printing, directing both filament geometry and cellular behavior. Support bath stiffness regulated mechano-sensitive gene expression and microtissue phenotype, with softer matrices favoring chondrogenesis and stiffer environments promoting (myo)fibrogenic differentiation. In addition, the physical properties of the support bath modulated microtissue fusion and extracellular matrix organization, with increased collagen fiber alignment in stiffer baths. Leveraging these findings, it was possible to engineer either articular cartilage, meniscus, or ligament grafts with user defined collagen architectures by simply varying the physical properties of the support bath. This platform establishes a foundation for bioprinting structurally anisotropic and phenotypically distinct constructs, thereby enabling the scalable engineering of a range of different musculoskeletal tissues.

## 1. Introduction

The field of tissue engineering (TE) seeks to regenerate tissues or organs impaired by trauma or disease. Achieving this goal will require the engineering of regenerative grafts that replicate the complex structure, composition, and function of the native tissue. This is particularly true for the repair of musculoskeletal tissues, such as articular cartilage (AC), meniscus, ligament and tendon, whose unique biomechanical function derives from the specialized architecture of their extracellular matrix (ECM). Damage to these tissues commonly initiates further joint degeneration, leading to the development of debilitating diseases such as osteoarthritis (OA) [1, 2]. Previous attempts at engineering such tissues have focused on seeding stem/progenitor cells within a 3D scaffold or a hydrogel, which are then stimulated in an attempt to promote tissue-specific ECM deposition [3–6]. However, these strategies typically fail to produce phenotypically defined tissues with biomimetic collagen network architectures that are integral to their physiological function, dramatically limiting their clinical utility.

Limitations associated with traditional TE strategies has motived increased interest in scaffold-free approaches for the development of regenerative grafts. These strategies harness the ability of cells to self-organize in response to cell-cell and cell-matrix interactions, mimicking the self-organization observed in polarized tissues during embryonic development and during adult tissue self-renewal [7]. Under specific physicochemical conditions, key aspects of such self-organization can be replicated *in vitro*, allowing stem cells to generate microtissues (μTs) or organoids that mimic key features of a specific tissue or organ [8]. Under the appropriate conditions, it may be possible to combine such biological building blocks to engineer scaled-up tissue grafts [9–12]. Realizing this goal will require the development of biofabrication strategies that can not only control the phenotype of the resulting graft, but also can also direct its (re)modelling to enable the engineering of tissues with user defined extracellular matrix architectures.

During development, individual tissues almost never develop in isolation, but do so concurrently with surrounding tissues and organs, which mechanically confine, impinge upon or pull on them [13, 14]. These mechanical influences have been postulated to affect the development of different tissues and indeed the entire early mouse embryo [15]. When cultured *in vitro* in the presence of physical boundaries, cells maintain the ability to sense and respond to such geometrical and physical cues [16]. For example, when endothelial cell sheets are cultured on substrates with distinct geometries (e.g., squares or annular rings), cells at the center stop proliferating upon reaching high density, whereas those at the periphery and at mechanically stressed regions (such as edges and corners) continue dividing. Similarly, the differentiation of adult mesenchymal stem/stromal cells (MSCs) is strongly influenced by geometric confinement and substrate stiffness [17–20], with stiffer substrates and increased cytoskeletal tension promoting an osteoblastic phenotype [21]. These geometric and mechanical signals introduce an additional layer to cell signaling, imparting information to individual cells and guiding the formation of fully developed tissues [22]. Therefore, by providing spatially and temporally defined physical boundary conditions to populations of microtissues or organoids, it may be possible to direct their fusion and (re)modelling in order to engineer scaled-up constructs that mimic the anatomical shape, internal structure, composition and function of specific tissues and organs.

3D bioprinting is now an established technique to spatially pattern cells and supporting biomaterials to replicate key anatomical features of native tissues. Such approaches have recently been extended to the bioprinting of cellular aggregates, microtissues and organoids [23–26]. However, current bioprinting approaches often struggle to achieve precise microtissue patterning and adequate fusion between microtissues [27], which is critical requirement for the engineering of scalable grafts. Moreover, little consideration has been given to how the bioprinting process can be used to direct long-term phenotype and structural (re)modelling of the resulting tissues. This could potentially be addressed using emerging 4D bioprinting concepts, and in particular the use of support baths that temporally adapt to provide precise biophysical cues to printed cells [28, 29]. Recognizing that normal tissue development depends on both the self-organizing potential of stem cells and key physicochemical cues from the microenvironment to establish its final architecture and function, here temporally adapting support baths will be used to control the surrounding matrix stiffness and the degree of physical confinement experienced by MSC derived microtissues post-printing. To this end, extrusion based bioprinting platform will be used to precisely pattern microtissues at a high density within a methacrylate xanthan gum (XG-MA) support bath. By precisely tuning the physical properties (rheology, stiffness) of the support bath, we demonstrate that is it not only possible to modulate print fidelity, but to also direct microtissue fusion, differentiation and the long-term structural organization of the bioprinted graft.

## 2. Results and discussion

### 2.1 Development of a temporally adapting support bath for extrusion bioprinting of microtissues

To facilitate the bioprinting of large numbers of microtissues into a support bath, we first needed to identify a sacrificial supporting bioink that would rapidly vacate the support bath to enable microtissue fusion post-printing. Gelatin has previously been used as a sacrificial ink in support baths, enabling the 3D printing of perfusable channels for vascular TE [30, 31]. Therefore, we first assessed the rheological properties of gelatin at a range of different concentrations (Figure S1A). While all gelatin concentrations displayed favorable shear thinning properties, we selected a relative low concentration (1% w/v) to minimize the shear stresses experienced by the microtissues during the bioprinting process. Another key advantage of gelatin is its temperature-dependent viscosity, displaying a higher viscosity at lower temperatures, thereby preventing settling of microtissues within the print cartridge if cooled to 4°C for 15 minutes before printing.

In order to generate a continuous filament of tissue, we next needed to determine how the distribution of microtissues post-bioprinting was influenced by their density within the supporting gelatin bioink. The ideal μT density for extrusion bioprinting will depend on the needle size and the μT dimensions, with high densities hindering extrusion [32] or imposing the cells to very high shear stresses, thereby negatively impacting viability [33, 34]. Here we found that microtissues measuring 175 ± 5 μm in diameter (Figure S1C) could be extruded at relatively high densities using a 22G metal needle (outer diameter of 0.7 mm and an inner diameter of 0.41 mm). Bioprinting at low μT densities (2,000 μT mL^−1^) resulted in clear gaps within the resulting filament (Figure 1B), while higher densities (60,000 μT mL^−1^) caused clogging during printing, which also resulted in the development of gaps (Figure 1B). We identified 45,000 μT mL^−1^ as the optimal density to form compact, continuous tissue filaments post-printing (Figure 1B).

**Figure 1.**
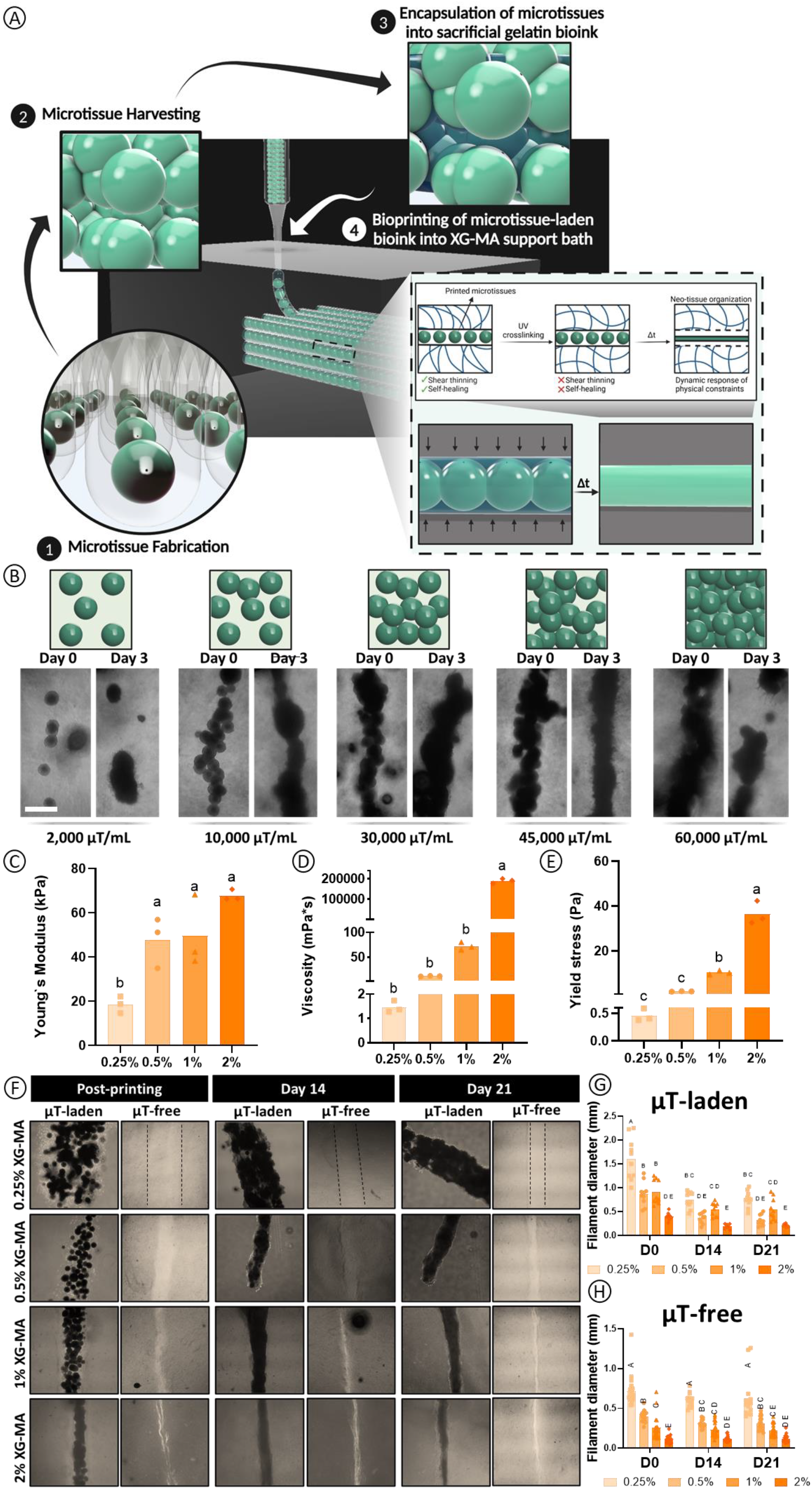
The concentration of the support bath influences the extent of filament confinement over time post-printing. A) Graphical abstract of the 4D-bioprinting platform. Microtissues are fabricated using a high throughout agarose microwell system and cultured for 2 days prior to their harvest and encapsulation into a gelatin bioink and extrusion into a XG-MA support bath. After UV-crosslinking, the filament diameter changes over time due to the changing physical confinement provided by the support bath. B) The influence of microtissues density on the development of a continuous tissue filament 3 days post-printing. C) Youngs Modulus of XG-MA hydrogels following UV crosslinking for 4 minutes. D) Viscosity of the un-crosslinked XG-MA bath at different concentrations. E) Yield stress of the un-crosslinked XG-MA bath. F) Brightfield microscope imaging of the printed microtissues within XG-MA support baths of differing concentrations at different time points. Filament diameter changes over time for the (G) μT-laden inks and the (H) μT-free inks. Scale bar (Sb): 200μm. 2-way ANOVA followed by the Tukey`s post comparison test was performed to assess the differences between the groups at different time points; significance was accepted when p<0.05. Statistically comparable groups are labeled using compact letter display where groups sharing the same letter are not significantly different (p>0.05).

To enable the high resolution bioprinting of μTs, we next sought to extrude them into a support bath with tailorable mechanical and rheological properties. We selected xanthan gum (XG) as a support bath and further modified it with a methacrylate group (XG-MA) to stabilize and mechanically support the printed filament throughout the culture process (Figure S1A). The reaction between glycidyl methacrylate and the hydroxyl or carboxyl groups of XG was confirmed through ^1^H-NMR analysis, where the presence of the two peaks at 6.58 and 6.17 ppm (Figure S1A) confirms the modification of the XG chain with a methacrylate (−C=C) group, enabling the material to be crosslinked post UV exposure. 1.5% XG or 1% XG-MA have previously been used as a support bath for bioprinting [35–36]. To expand on this, we tested a range of XG-MA concentrations (from 0.25 to 2% w/v) and measured their stiffness post UV crosslinking. We found that the Young’s modulus increased with XG-MA concentration, from 20 kPa for the softest bath to 65 kPa for the stiffest bath (**p < 0.01; Figure 1C).

We also investigated the shear thinning and thixotropic properties of the support baths to assess their suitability for bioprinting applications (Figure S2). All tested concentrations of XG-MA displayed shear-thinning (Figure S2A, B) and thixotropic (Figure S2C) properties, allowing the bath to maintain its structural integrity post-printing. Irrespective of concentration, the XG-MA was able to recover after experiencing high shear stress, with a faster recovery observed in the higher concentration baths (Figure. S2C). Having developed a support bath with tunable mechanical and rheological properties, we next sought to assess its capacity to support high resolution extrusion bioprinting of high density microtissues-laden bioinks. We first sought to assess how the concentration of XG-MA with the support bath influenced the geometry of the resultant print following extrusion bioprinting of a microtissue-laden gelatin ink and a microtissue-free gelatin ink. Using the same extrusion printing parameters (6 uL s^−1^ extrusion rate, 2 mm s^−1^ printing speed), a higher concentration XG-MA support bath enabled a higher printing fidelity/resolution (i.e. a smaller printed filament diameter) for both a μT-free and a μT-laden ink (Figure 1G, H). The printed microtissues generated a larger diameter filament when printed into a softer bath, ranging from 1.5-2mm in diameter following extrusion into a low stiffness 0.25% bath, to 0.25-0.5 mm in the stiffer 2% bath for both μT-laden (Figure 1G) and μT-free inks (Figure 1H). 7 days post-printing, the μTs appeared to have fused, with high cell viability observed in all support baths (Figure. S3A). This suggests that the shear stress applied to the cells within the μTs during the bioprinting process, as well as UV exposure to crosslink the support bath post-printing, does not negatively impact cell viability.

Not only did the concentration of XG-MA in the support bath influence the initial diameter of the printed filament, but it also impacted how it changed with time in culture. In μT-laden bioinks, the filament diameter was found to reduce in diameter with time in culture, with larger absolute reductions observed in the softer baths (Figure 1F, G). This is likely due to both cellular remodeling and physical forces exerted by the support bath on the printed μTs as the supporting gelatin ink washes out with time. However, even in the μT-free prints (where no cellular remodeling is involved), a reduction in filament diameter over time is observed, pointing to an active role for the support bath in determining the long-term geometry of the print (Figure 1F, H). Collectively these results suggest that the support bath exerts a radially compressive force on the filament post-printing, resulting in a temporal reduction in diameter until an equilibrium is reached between day 14 and 21 (Figure 1H).

These results suggest that the XG-MA support bath can be used to support the 4D bioprinting of μTs, where the printed structures will change in shape, function and/or behavior over time in response to external stimuli provided by the bath. In traditional 4D printing approaches, this dynamic transformation is typically achieved using materials that respond to environmental factors such as temperature, pH, humidity etc. [37–39]. For 4D bioprinting, this has commonly involved the supporting material ink changing over time (e.g. shrinking), which in turn influences the final tissue shape and organization. Here, is not the bioink (gelatin) that changes over time, rather the physical boundaries that are provided to the printed μTs in the form of the temporally changing support bath. Previous studies have shown that physical boundaries can guide neo-tissue organization, enabling the engineering of anisotropic living systems [40, 41]. Furthermore, tissue maturation *in vivo* is driven by spatiotemporal cellular signaling and mechanical forces [42]. Therefore, we hypothesized that the degree of physical confinement imposed by the bath over time can influence microtissues fusion and phenotype, as well as the structural organization of the resulting tissue, thereby enabling the bioprinting of anisotropic grafts.

### 2.2 YAP activation in MSC derived microtissues is influenced by support bath stiffness

MSCs are known to respond to the stiffness of their surrounding environment [43, 44]. As the stiffness of the support bath increased with XG-MA concentration (Figure 1C), we next investigated whether such differences in the local mechanical environment would influence the activity of known mechanosensitive proteins in the bioprinted microtissues (Figure 2A). The activation of YAP, a key protein in the Hippo signaling pathway that regulates growth and development, is known to be influenced by mechanical cues in the ECM for diverse cell types, including cardiac cells, myofibroblasts and MSCs [45–50]. YAP is regulated by actomyosin cytoskeletal tension and functions as a transducer of mechanical cues, with increased ECM stiffness and cell spreading leading to higher YAP activity [22]. This in turn can influence cellular differentiation, with YAP activation known to promote osteogenesis [51], while suppressing adipogenic [52] and chondrogenic differentiation [50]. Here, bioprinted microtissues were maintained in media supplemented with TGF-β3 post printing, a growth factor known to support either a chondrogenic or a myogenic phenotype in MSCs depending on other environmental factors such as the ability of cells to spread [43, 53–55]. To assess YAP activity across different bioprinting conditions, we performed YAP staining at both early and late time points in constructs printed into XG-MA support baths of differing stiffness (Figure 2B). YAP activity was significantly higher in microtissues printed into the stiffer support baths (1% and 2% XG-MA). In these stiffer conditions, greater YAP nuclear colocalization was also observed (Figure 2A, white arrows). This suggests that the greater stiffness of higher concentration XG-MA baths bath promotes YAP transcription and its activation.

**Figure 2.**
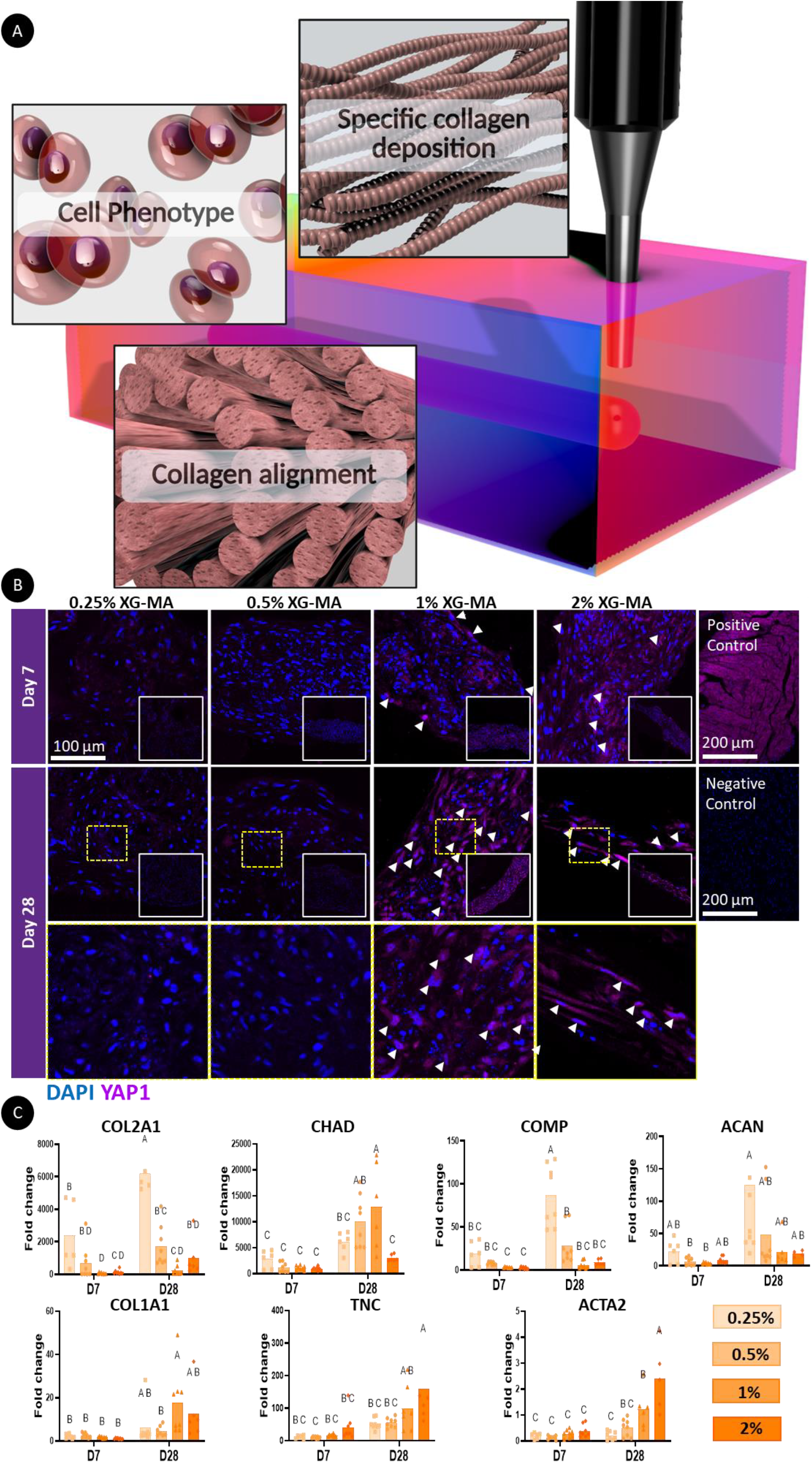
The physical properties of the support bath influence MSCs phenotype. A) Graphical abstract of a 4D bioprinting platform capable of generating phenotypically distinct musculoskeletal grafts. B) YAP1 staining of printed microtissues reveals spatial differences in YAP expression depending on the stiffness of the support bath. White arrows indicate regions where YAP colocalizes with the nuclei, indicating YAP activation. Positive control is mouse heart and negative control is mouse heart without primary antibody. C) Expression of a range of genes associated with a cartilage (COL2A1, CHAD, COMP, ACAN), fibrocartilage or ligament tissue (COL1A1, TNC, ACTA2). For statistical analysis, a two-way ANOVA test was performed followed by Tukey’s post comparison to compare the means of each group at different time points. Significance was accepted when p<0.05. Statistically comparable groups are labeled using compact letter display where groups sharing the same letter are not significantly different (p>0.05).

### 3.3 Support bath stiffness regulates microtissue phenotype

It is well established that physical and biochemical signals work in synergy to guide cell fate and function [56, 57]. As YAP activity within the bioprinted microtissues was strongly regulated by the stiffness of the surrounding support bath, we further hypothesized that longer-term MSC phenotype would also depend on these physical cues. To this end, the printed constructs were cultured in the presence of TGF-β3 and MSC phenotype was assessed over a 4-week culture period using a range of biochemical and histological assays. We first assessed the expression of a range of genes associated with chondrocytes, fibrochondrocytes, and myofibroblasts (Figure 2C). Actins are major cytoskeletal proteins involved in cell migration and contraction. Alpha smooth muscle actin (α-SMA) is particularly associated with contractile fibroblastic cells such as myofibroblasts, which play a role in tendon and ligament healing and maintaining tissue homeostasis [58]. The highest levels of ACTA2 expression (a gene that encodes for smooth muscle alpha-2 actin) was observed in the stiffer 2% baths after 28 days of culture (Figure 2C). We also assessed α-SMA deposition through immunofluorescence staining (Figure 3A). While weak α-SMA staining were observed at day 7 (Figure 3A), a significant increase was evident by day 28, particularly in the stiffer (1% and 2%) baths (p<0.05) (Figure 3A, B). COL1A1 and TNC (Tenascin-C) gene expression, markers associated with a ligamentous or tendon phenotype, was also higher in the stiffer baths (Figure 2C). In agreement, type I collagen deposition was higher in the stiffer support baths at both early and late time points (Figure 3A, C).

**Figure 3.**
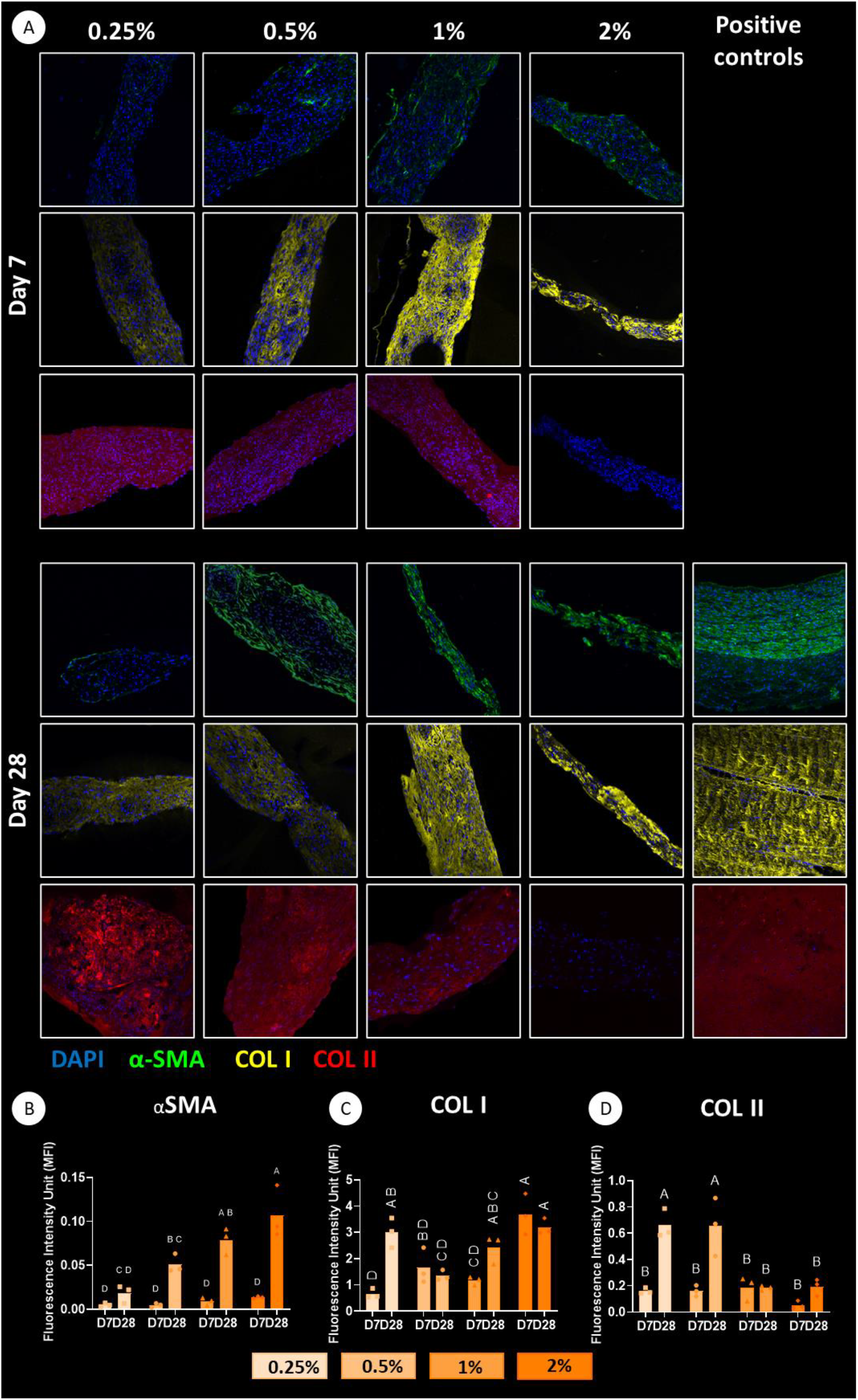
Immunofluorescence staining for specific proteins within the bioprinted microtissues. (A) Staining for type I collagen, type II collagen and α-SMA deposition in 7 and 28 days post printing in the different support baths. Positive controls are arterial control for α-SMA, ligament for type I collagen and articular cartilage for type II collagen. Quantification of immunofluorescence intensity normalized with background intensity and normalized to the number of nuclei of each region of interest (ROI) for αSMA (B), collagen I (C), collagen II (D); n=3. For all the graphs a two-way ANOVA test was performed followed by Tukey’s post comparison to compare the means of each group at different time points. Significance was accepted when p<0.05. Statistically comparable groups are labeled using compact letter display where groups sharing the same letter are not significantly different (p>0.05).

While genes and proteins associated with a ligament or tendon phenotype were more highly expressed in the stiffer bath, the expression of the chondrogenic genes COL2A1, COMP and ACAN were higher in microtissues bioprinted into the softer support baths (Figure 2C). In agreement with this finding, strong staining for type II collagen deposition was observed in the tissues generated in the soft baths, with little deposition observed in the stiffer baths (Figure 3A, D). Together these findings suggest that the MSC derived microtissues actively respond to the mechanical cues provided by the different support baths. The softer 0.25% and 0.5% XG-MA support baths promote greater collagen II deposition and reduced alpha-smooth muscle actin (ACTA2) expression, resulting in a more chondrogenic phenotype (Figure 3). The observation of enhanced sGAG accumulation in these softer environments further confirms this conclusion (Figure S3B). The 1% XG-MA bath, which possess an intermediate stiffness, appeared to support a more fibrochondrogenic phenotype, as evident by simultaneous sGAG (Figure S3B) and type I collagen deposition (Figure 3A, C). Finally, the stiffest 2% XG-MA baths supported the highest levels of type I collagen production (Figure 3A, B) but little sGAG deposition (Figure S3B), suggesting the development of a more ligamentous tissue.

### 3.4 Support bath stiffness regulates collagen fiber alignment in bioprinted tissues

Encouraged by the successful bioprinting of high density microtissue-laden bioinks into a temporally adapting support bath, where the microtissues fused effectively over time and adopted a phenotype defined by the stiffness of the surrounding support bath, we next sought to explore if this bioprinting platform could be used to direct collagen organization and hence enable the engineering of anisotropic soft tissues. The bioprinting of such structurally organized tissues has previously been demonstrated using collagen-based bioinks, where the extrusion process directed collagen fiber alignment within the inks, which in turn facilitated cell spreading and alignment along the fiber direction, thereby promoting the formation of anisotropic tissue [59–61]. However, the slow gelation and relatively poor printability of collagen-based bioinks challenge this approach [62, 63]. Here we sought to guide collagen alignment by printing microtissues into a physically constrained environment that facilitates microtissue fusion and directs subsequent ECM organization. To validate our ability to bioprint custom-shaped, highly aligned musculoskeletal tissues, we employed polarized light microscopy (PLM) to assess the preferential alignment of collagen fibers along the printed filaments (Figure 4). After four weeks of *in vitro* culture, the microtissues fused to generate filaments of tissue whose internal structure was directed by the physical constraints imposed by the XG-MA support bath. Picrosirius red (PR) staining (Figure 4A) revealed robust collagen deposition across all groups, with more homogeneous collagen distribution observed within filaments printed in the stiffer baths (Figure 4A). Collagen deposition, quantified relative to DNA levels (Figure 4E), was also significantly higher in the stiffer baths (p<0.01). In addition, the XG-MA bath provided effective boundary conditions for directing collagen organization in the neotissue. While collagen fibers tended to align parallel to the print direction (i.e. at 0 °C) irrespective of the bath stiffness (Figure 2B), the stiffer 2% support bath supported the greatest degree of fiber alignment (Figure 2A) and coherency (Figure 2D). We also investigated the size of the collagen fibrils through SEM imaging (Figure S4), where preferential alignment of the collagen fibers was observed on the surface of all groups. The diameter of the collagen fibril ranged between 20 and 60 nm (Figure 4C) after 4 weeks of *in vitro* culture, which is comparable to that observed in the cartilage of skeletally immature pigs [64].

**Figure 4.**
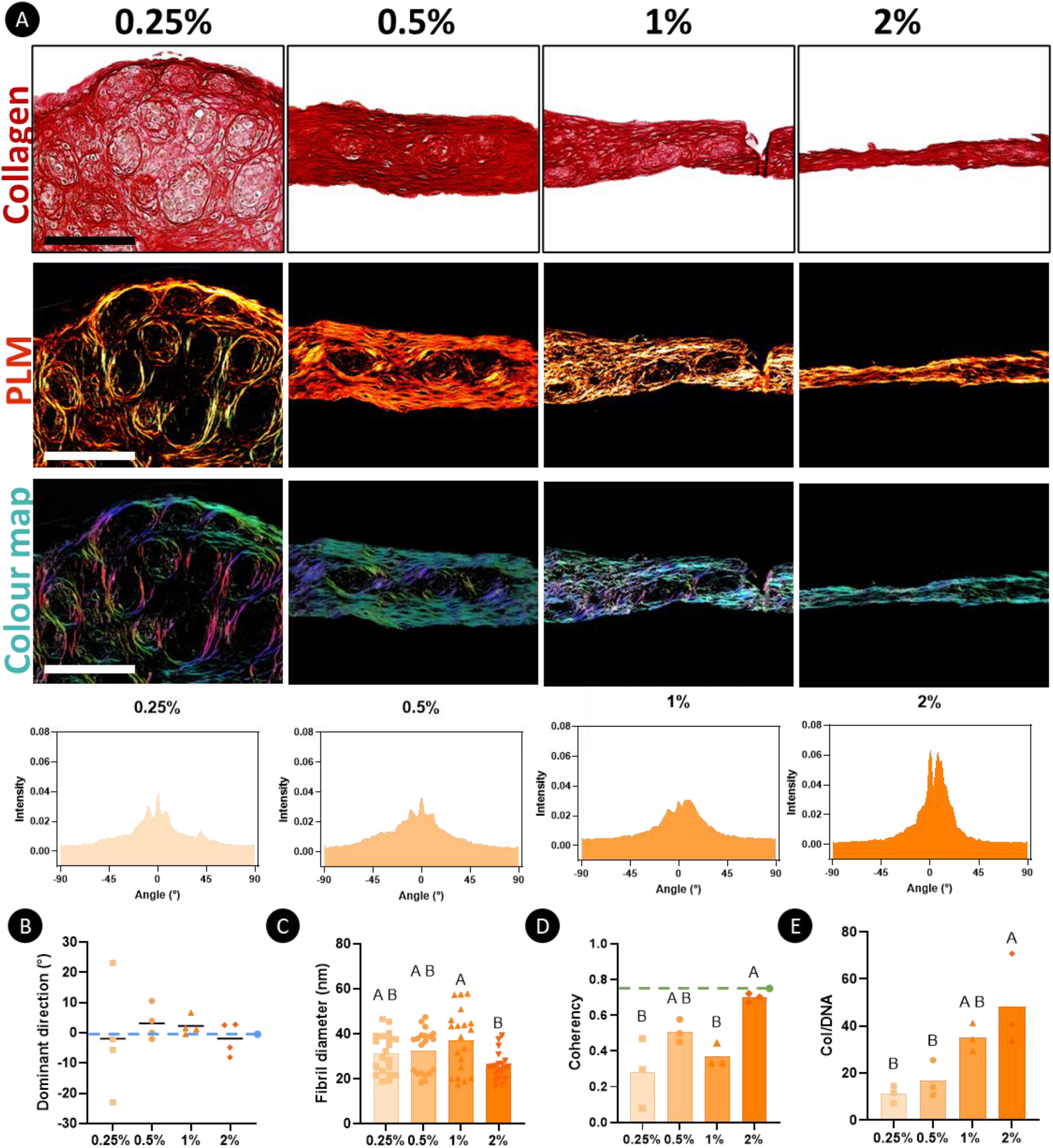
A stiffer support bath improves microtissue fusion and the alignment of the secreted collagen fibers. A) Histological assessment of bioprinted microtissues after 28 days post printing with picrosirius red staining, using PLM imaging to analyze collagen fiber alignment. Color maps are generated from PLM images. Here, color hue is used to indicate fiber orientation, where blue/cyan indicated fibers oriented at 0 degrees and pink/red indicated fibers oriented at 90 degrees. Sb: 200μm. Quantification of the fiber orientation was then performed for the 0.25%, 0.5%, 1% and 2% XG-MA support baths. B) Average collagen fiber orientation with the filaments generated in the different support baths. C) Fibril diameter at 28 days post printing. D) Fiber coherency, where values close to 1 indicated that fibers are aligned in the same direction, while a value closer to 0 indicated a higher dispersion of the fibers in different directions (n= 4). E) Biochemical quantification of total collagen normalized to DNA levels. Statistical difference is determined using a two-way ANOVA test with Tukey’s post comparison test. Significance was accepted with p<0.05. Statistically comparable groups are labeled using compact letter display where groups sharing the same letter are not significantly different (p>0.05).

### 3.5. Bioprinting of anisotropic musculoskeletal tissues by spatial patterning microtissues into support baths to differing stiffness

Musculoskeletal tissues, particularly those that endure mechanical loads in specific directions, exhibit preferential collagen fiber alignment to support their biomechanical functions. For example, tendons and ligaments have significantly higher tensile properties along the fiber direction [65]. In the meniscus, the predominance of circumferentially aligned type I collagen fibers provides the necessary stiffness (100–300 MPa) to enable this tissue to function within the knee joint [66]. Similarly, in AC vertically oriented collagen fibrils in the deep zone protect the tissue against large compressive strains [67], while collagen fibers aligned parallel to the surface in the superficial zone play a crucial role in resisting high tensile stresses [68, 69]. To demonstrate the utility of this microtissue based bioprinting platform, we next sought to engineer larger tissue grafts with distinct phenotypes and anisotropic collagen fiber networks. As a proof of concept, we sought to replicate the distinct collagen arrangements observed in different musculoskeletal tissues, specifically the arcade-like collagen alignment in AC [69, 70], circumferentially aligned fibers in the menisci [71], and the linear collagen orientation found in ligaments [72]. To maintain anisotropy across multiple layers of printed microtissue filaments, we optimized filament spacing within the support bath to promote tissue compaction while preserving the desired collagen organization (Figure S5). For all of these prints, the composition of the culture media remained constant (a chemically defined media supplemented with TGF-β3), with only the stiffness of the bath changing for the bioprinting of the different tissue types.

Microtissues were first extruded into a 0.5% XG-MA support bath to generate an arcade-like geometry characteristic of native AC, where they fused and reorganized over time, aligning according to the print path, and generating a cartilaginous tissue rich in type II collagen (Figure 5D). In the superficial zone of the resultant tissue, collagen fibers predominantly aligned parallel (0 °C) to the surface of the construct, resembling that observed in native AC (Figure 5A (I, IV)). In the deep zone, fibers were primarily oriented at 90 °C, reflecting the perpendicular alignment seen in native AC (Figure 5(III, IV)). PLM quantification confirmed the superior tissue organization resulting from the support bath’s temporal confinement, leading to effective microtissue fusion and ECM reorganization (Figure 5A). Furthermore, the soft XG-MA support bath supported a chondrogenic phenotype, characterized by robust deposition of the hyaline cartilage specific matrix markers sGAGs and type II collagen (Figure 5D). For the meniscus construct, microtissues were bioprinted in the intermediate stiffness 1% XG-MA support baths in a circumferentially aligned pattern to replicate the organization seen in the meniscal ECM. PLM analysis confirmed that the final construct displayed high collagen fiber anisotropy, with fiber orientation closely matching the printing direction (Figure 5B). The meniscus is composed of two distinct regions: the inner zone, which contains a mix of collagen types I and II, and the outer zone, primarily made up of collagen type I. Here we successfully engineered a highly anisotropic meniscal tissue with uniform sGAG deposition and a high type I collagen content (Figure 5E)—characteristics typical of the outer zone, which accounts for over 80% of the meniscus body [73]. As a final proof of principle, we attempted to bioprint a ligament-like tissue within the stiffer 2% XG-MA support bath (Figure 5C, F). To this end, we bioprinted a three-layered structure with parallel collagen alignment to replicate the ligament architecture (Figure 5C). Our results demonstrated that MSC derived microtissues adapted to their surroundings, secreting type I collagen and expressing α-SMA (Figure 5 F), indicating (myo)fibroblastic differentiation [74]. Collectively, our results highlight that the physical boundaries provided by the support bath, applied to microtissue-rich bioinks, can be used to direct neo-tissue organization and enable the engineering of phenotypically distinct musculoskeletal tissues.

**Figure 5.**
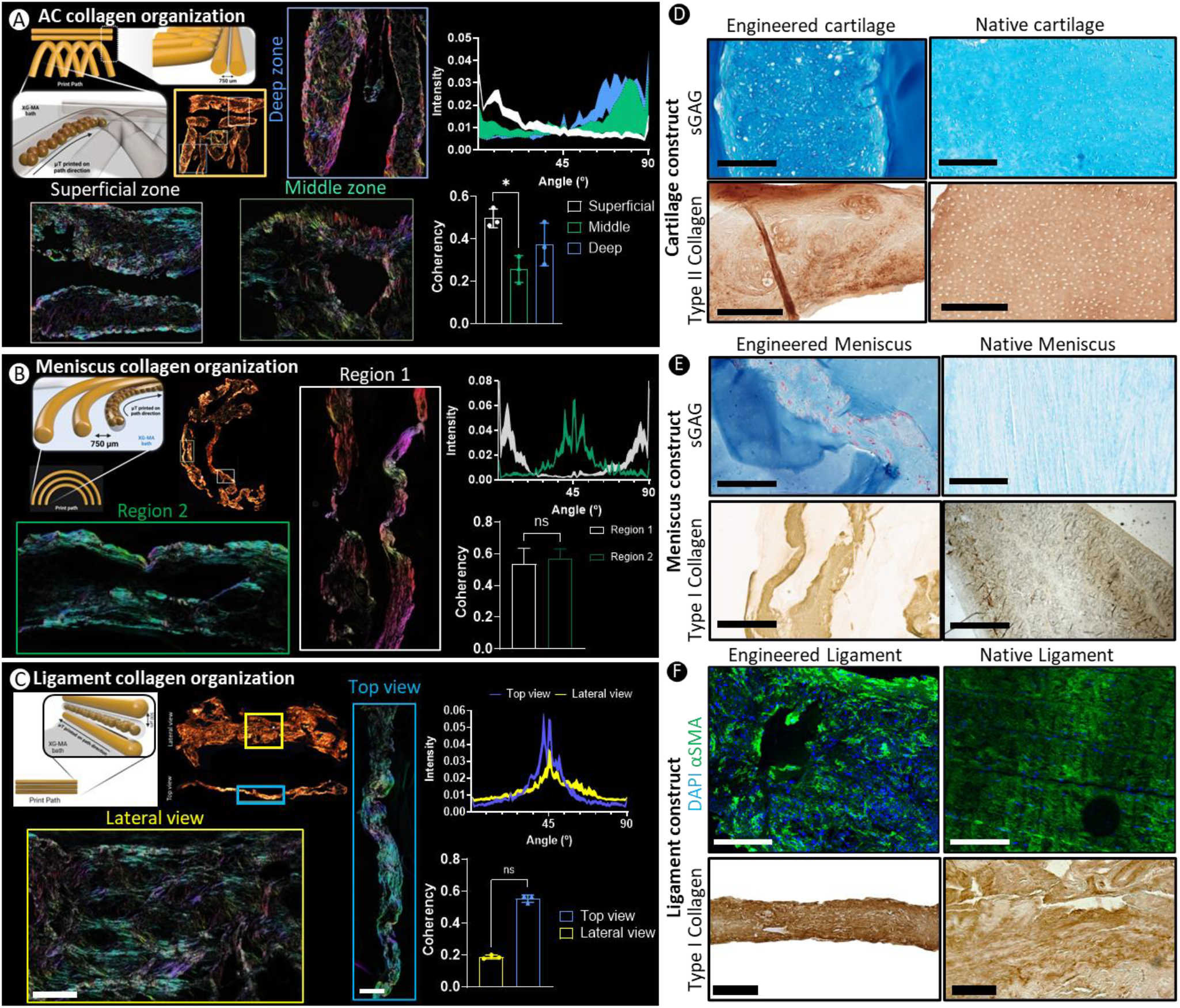
The bioprinting of scaled-up anisotropic tissue grafts. A) Bioprinting of Benninghoff collagen arcades typical of native AC. The collagen fiber orientation was quantified in the superficial, middle and deep regions of the tissue, indicating the development of highly anisotropic tissues. Coherency analysis values closer to 1 indicate that the fibers are aligned in one direction, while values close to 0 indicates the fibers are aligned in different directions. White indicates the superficial zone, green indicates the middle zone, blue indicates the deep zone. B) Bioprinting of circumferential filaments to engineer a meniscal tissue. Images from two different regions (white and green box) are provided, again demonstrating that the collagen fibers are aligning according to the printing path. Fiber coherency analysis indicates that most of the fibers are aligned in one direction. The white lines/bars represent the collagen fiber orientation within the regions highlighted within the white box (Region 1). The green lines/bars represent the collagen fiber orientation within the regions highlighted within the green boxes (Region 2). C) Bioprinting of a 3-layered anisotropic ligament tissue, where most of the fibers are aligned towards the 0 °C direction in both the lateral (yellow color box) and top (blue color box) view. A one-way ANOVA test was performed followed by Tukey’s post comparison to compare the means of each group at different time points. Significance was accepted when p<0.05. * Indicates p<0.05. ns: non-significant. For all the graphs: n=3. D) Histological and immunohistochemical assessment for the cartilage construct for sGAG and type II collagen deposition and compared to native AC. E) Histological and immuno-histological assessment for the meniscal construct for sGAG and type I collagen and compared to the native menisci. F) Histological and immunohistochemical assessment for the ligament construct for α-SMA and type I collagen. Scale bars: 200μm.

## 3. Conclusion

In summary, this novel bioprinting platform, which leverages a temporally adapting support bath, creates a dynamic environment that enables the precise spatial patterning of microtissue-laden bioinks. We demonstrate that the physical environment created by the surrounding support bath can modulate the phenotype of the printed microtissues, enabling the engineering of different musculoskeletal tissues by simply varying the support bath stiffness. Remarkably, this approach not only supports distinct musculoskeletal phenotypes, but also controls microtissue (re)modelling and subsequent ECM organization, thereby enabling the engineering of anisotropic grafts with user-defined collagen architectures. Furthermore, we demonstrate the versatility of this platform by bioprinting a range of different musculoskeletal tissues, including articular cartilage, meniscus, and ligament grafts. These findings support the continued development of 4D bioprinting strategies that more closely mimic normal developmental processes to better engineer functional tissues or organs. In the future such bioprinting platforms could potentially provide implantable, functional grafts at scale for a range of different regenerative medicine applications.

## 4. Materials and Methods

### 4.1 Modification of Xanthan Gum with methacrylic group (^1^H-NMR) and support bath preparation

Xanthan gum (XG) (Sigma) was dissolved at 0.5% (w/v) into ultra-pure water and stirred until completely dissolved. Once dissolved, glycidyl methacrylate (40 mL) were added to the XG solution and stirred overnight at 60 °C, protected from light. The solution was now dialyzed for 7 days against DI with water changes twice a day, freeze dried for 48 hours and stored at −20 °C for long term storage. To ensure sterilization, aliquots of XG-MA were weighted and placed into a 50 mL tube and sterilized in an ethylene oxide gas sterilizer (Andersen Products) for 12 hours. 2 days prior printing, the material was dissolved into phenol free DMEM (Gibco) supplemented with penicillin (100 U mL^−1^), streptomycin (100 μg mL^−1^) (P/S) (both Gibco) and let it rotate at 40 rpm at RT for 2 days. The day of printing, Lithium phenyl-2,4,6-trimethylbenzoylphosphinate (LAP) (Sigma), was added for a final concentration of 0.25% (w/v) into the XG-MA solution and vortexed. To ensure the removal of the bubbles, the bath was centrifuged at 2500 x *g* for 5 minutes at RT, prior printing. For each print, XG-MA (4 mL) was pipetted into a 6 well plate with a pipette for viscos liquids (Microman E, Gilson) to ensure no formation of bubbles during the pipetting.

### 4.2 Rheological assessment of bioink and support baths

Rheological characterizations were conducted on an MCR 102 Rheometer (Anton-Paar, Hertford Herts, UK) equipped with a Peltier element for temperature control. A plate-plate geometry with a diameter of 25 mm (PP25) was used in all the tests. The viscosity as a function of shear rate (0.1–1000 s−1) was conducted at a constant temperature of 4 °C for the gelatin (Gelatin Type B) (Sigma) bioink, and at RT for the XG-MA support bath. To investigate the linear viscoelastic region (LVR) and determine the yield stress, an amplitude sweep test was carried out at RT. An angular frequency between 1 and 100 rad s^−1^ was applied to the sample to determine G′ and G″ curves. The G′ and G″ crossover point was taken as the yield point where the material behavior transition from solid to liquid-like behavior [75]. A dynamic sweep stress test is performed at a shear strain range of 1.0 to 100% to determine the solid and liquid-like state of different bio-inks. Five intervals at a constant shear rate were performed to determine the thixotropy behavior of the support bath and investigate the time dependency of the support bath. Here, the viscosity was measured in transient shear rate step tests at RT. First, samples were sheared at a shear rate of 0.03 s^−1^ for 30 s, then the shear rate increased to 10 s^−1^ for 30 s, and again, the shear rate decreased to 0.03 s^−1^ for 30 s. Bioinks and support baths were kept in a high humidity atmosphere to prevent dehydration from affecting the rheological results. All measurements were performed in triplicate.

### 4.3 Compression testing of XG-MA hydrogels

Unconfined <u>compression tests</u> were carried out on samples produced in the shape of a cylinder using a 5 mm PDMS mold and crosslinked with UV for 4 minutes in the mold. Samples were taken out of the mold and were placed in a PBS bath to prevent the dehydration, and therefore compressed at a rate of 5 mm s^−1^ using a twin column Zwick universal testing machine (Zwick, Roell) equipped with a 10 N load cell. A <u>preload</u> of 0.01 N was used for the hydrogels. Test was stopped when the gel underwent breakage. The Young’s modulus was defined as the slope of the linear phase of the resulting stress-strain curve of the compression to 70% strain.

### 4.4 Isolation and expansion of caprine BM-MSCs

Caprine bone marrow mesenchymal stem cells (MSCs) were isolated from the sternum of skeletally mature, female goats. The bone marrow was cut into 2mm^3^ pieces, vortexed for 2 minutes in high glucose DMEM (hgDMEM) (Gibco) supplemented with P/S, 10% Fetal Bovine Serum (FBS) (Gibco) (Expansion medium, XPAN) to aid liberate cellular components and then seeded at a density of 5.7×10^4^ cells cm^−2^ and maintained in a humidified atmosphere under physioxic conditions (5% CO_2_, 5% O_2_). After 72 hours, the flasks were washed twice with sterile PBS to remove non-adherent cells and debris. Cell culture media was exchanged to fresh XPAN every 2 days. When the MSCs colonies were 80% confluent, cells were frozen and stored in liquid nitrogen for long term storage. For further expansion, cells were seeded at a density of 5 × 10^3^ cells cm^−2^ at 5% O_2_ and expanded up P3 for all bioprinting experiments.

### 4.5 Fabrication of MSC microtissues (μTs)

The μT were fabricated as previously described [76]. The 3D printed mold stamps containing 1,889 micro resections and 4% (w/v) ultrapure agarose were sterilized in autoclave at 120 °C for 20 minutes. Next, ultrapure agarose (6 mL) (Sigma) was gently poured into each well of 6-well plates and allowed to cool down for 15 min after molding it. Agarose molds were then soaked overnight in XPAN, at 37 °C in a humidified atmosphere with 5% O_2_. MSCs were seeded in a density of 7.556×10^5^ mL^−1^ into the microwells (5 mL), to obtain 1889 μTs with ∼2,000 cells/μT and centrifuges at 700 x *g* to collect the cells at the bottom of each well. The MSCs were maintained in chondrogenic media (CDM) consisting of hgDMEM GlutaMAX supplemented with P/S, sodium pyruvate (100 μg mL^−1^), L-proline (40 μg mL^−1^), L-ascorbic acid-2-phosphate (50 μg mL^−1^), linoleic acid (4.7 μg mL^−1^), bovine serum albumin (1.5 mg mL^−1^), 1× insulin−transferrin−selenium (ITS), dexamethasone (100 nM) (all from Sigma), and human transforming growth factor-beta 3 (TGF-β3) (10 ng mL^−1^) (Peprotech, UK). After 2 days, the μTs were harvested prior bioprinting. For this, media was first flushed over the agarose surface using a micropipette, and subsequently collected into a 50 mL tube as described previously [11]. The density of the μT suspension was quantified, and the appropriate volume was used to seed a precise number of μTs into each construct.

### 4.6 Bioink preparation and bioprinting parameters

Sterile gelatin type B (1% w/v) was dissolved in hgDMEM for 30 minutes at 37 °C in a water bath the day prior printing. After 30 minutes, gelatin is removed from water bath and placed at 4 °C overnight for gelation. μTs are harvested from the agarose microwells according to methods described before [11], placed in a 50 mL tube with media, the tube is centrifuged at 650 x *g* for 5 minutes at RT to collect all the μTs at the bottom of the tube. Gelatin was then added to reach desired microtissues density for bioprinting. Afterwards, the bioink is carefully loaded into the cartridge and placed into the fridge for 15 minutes prior bioprinting, in sterile conditions. The bioprinting is carried out using a syringe pump printhead (Cellink, Sweden), with extrusion rate of 6 uL s^−1^ and print speed of 2 mm s^−1^.

### 4.7 Live/Dead assay

Cell viability was assessed using a live/dead assay kit (Invitrogen, Bioscience). The constructs were first rinsed in PBS 1X and then incubated for 1 hour in a solution containing calcein (2 μM) and ethidium homodimer-1 (EthD-1) (4 μM), at 37°C. Following the incubation period, the constructs were rinsed 3X with PBS to remove any excess of the dyes and were imaged using a Leica SP8 scanning confocal microscope. Excitation wavelengths of 485 nm and 530 nm were used for calcein, while emission wavelengths of 530 nm and 645 nm were used for EthD-1. Maximum projection z-stack reconstructions were generated to analyze cell viability throughout the depth of the tissue, with the view from the top.

### 4.8 Scanning Electron Microscopy (SEM) imaging and analysis

Samples were fixed in 3% <u>glutaraldehyde</u> in 0.1 M <u>cacodylate</u> buffer (all from Sigma-Aldrich) at 4 °C for a minimum of 12 h. The samples were enzymatically treated according to methods described before (Gannon et al, 2015) to remove any excess of GAGs that could disrupt the imaging of the collagen fibers. They were then washed with 1X PBS and dehydrated in graded ethanol baths series (50-100% EtOH) and dried using critical point drying method to better preserve the surface structure of the biological sample. Samples were imaged with a Zeiss ULTRA plus <u>SEM.</u>

### 4.9 Histological Evaluation

Briefly, samples were fixed with 4% paraformaldehyde (PFA) solution overnight at 4°C. Following fixation, the samples underwent dehydration using a series of ethanol solutions (from 50% to 100% v/v), followed by clearing in xylene and embedding in paraffin wax (Sigma). The samples were sliced into 5μm tissue sections and rehydrated. Haematoxylin and Eosin (H&E) staining (Sigma) was used to assess the tissue morphology, the ECM, and the nuclei of the cells. 1% (w/v) solution of Alcian Blue (AB) 8GX in 0.1 M hydrochloric acid (HCl) was used to stain sGAG content and counter-stained with 0.1% (w/v) nuclear fast red to determine cellular distribution. A 0.1% (w/v) Picrosirius red (PR) solution was used to visualize collagen deposition (all from Sigma). Stained sections were mounted using Pertex (Avantor) solution and imaged using an Aperio ScanScope slide scanner (Leica, Germany).

### 4.10 Immunofluorescence staining and quantification

Immunofuorescence was performed for collagen type I (Abcam ab90395 1:400), collagen type II (Santa Cruz sc52658 1:400) as previously described [77]. For α-sma (Abcam ab124964 1:200) and YAP (Abcam ab3472 1:200) staining, wax embedded sections were rehydrated and washed 3X with 1X PBS. Antigen retrieval was performed in a sodium citrate buffer (10 mM), 0.05% Tween20 pH 6 (all from Sigma) in a humidified environment at 80 °C for 20 minutes. The slides were then washed with 1X PBS + 0.05% Tween20 (WB) supplemented with 10% donkey serum (Blocking buffer, BB) to allow the blocking of non-specific sites. Samples were then incubated with the primary antibody in the BB in a humidified chamber overnight at 4°C. Samples were then washed with WB and incubated with the secondary antibody (1:200) in BB for 1h at RT in the dark. Samples were then washed with WB and mounted with counterstaining mounting media supplemented with DAPI (Fluoroshield, Sigma). Prior to imaging the slides were sealed with nail polish and stored in the dark at 4°C. For immunofluorescence quantification, the intensity of 3 specific region of interests (ROI) within each sample-from 3 different replicates-was measured and quantified through ImageJ software. The intensity was then normalized by the number of nuclei in the ROI.

### 4.11 RNA isolation and quantitative Real-Time PCR

At 7 and 28 days after bioprinting of the μTs, the samples were washed 3X with 1X PBS and then snap frozen for further processing. RNA was isolated from the samples using the Trizol method (Sigma-Aldrich), following the manufacturer’s instructions. Chloroform extraction was performed, and the RNA was re-suspended in RNase-free water. The RNA samples were then stored at −80°C. For cDNA synthesis, a HighCapacity RNA-to-cDNA™ Kit (Applied Biosystems™), was used to transcribe 500 ng of RNA from each sample into cDNA (20 μL) via polymerase chain reaction (PCR), according to manufacturer`s instruction (37°C for 1 h, 95 °C for 5 minutes and 4°C for 5 minutes). Real-time PCR was performed using an Applied Biosystems instrument, and Taqman PCR Fast-Advanced master mix (ThermoFisher) was used for the PCR reaction. Goat-specific TaqMan probes (ThermoFisher) were used for gene amplification and 18S rRNA TaqMan probe was used as housekeeping gene for each well. The qPCR amplification followed the following cycles: 50°C for 2 minutes, 95°C for 10 minutes and 40 cycles at 60°C for 1 minute each. The levels of gene expression were quantified using the real-time PCR data and analyzed with the 2^(−ΔΔCt)^ method. The expression levels were normalized to the average expression of the 18S rRNA housekeeping gene and normalized with MSCs monolayer cells.

### 4.12 Filament diameter measurements and polarized light microscopy analysis

Diameter measurement of bioprinted μTs were conducted using a 4X objective of an inverted microscope (Primo Vert, Zeiss, USA), and calculated using ImageJ software (Wayne Rasband and Contributors, USA). Images were captured at different intervals until 28 days post printing. Four independent measurements of each construct were obtained from each experimental replicate.

For PLM analysis, sections were stained with 0.1% (w/v) PR, mounted with Pertex (all from Sigma-Aldrich) and imaged with an Olympus BX41 polarizing <u>light microscope</u> equipped with a MicroPublisher 6™ CCD camera and an Olympus U-CMAD3 adaptor. Average orientation, dispersion and <u>coherency</u> of collagen fibrils in the engineered tissues were assessed using orientationJ and <u>directionality</u> plugins in ImageJ.

### 4.13 Statistical analysis and graphical results

Statistical analyses were performed using the software package GraphPad Prism (version 10.0). Statistical tests used to assess the <u>normal distribution</u> of data or to compare groups are indicated in figure legends. When groups were compared, significance was accepted at a level of p < 0.05. Results are expressed as mean ± standard deviation. <u>Graphical results</u> were produced with GraphPad Prism.

## Supporting information

Supportin Information

## 6. Acknowledgements

We would like to thank Dr Megan Canavan from Trinity College Dublin for the SEM imaging. Schematic diagrams of graphical abstract, figures 1 and 2 were created with BioRender.com and Autodesk Fusion 360 software by F.D.S. This work was founded by the European Research Council (ERC, 4D-BOUNDARIES #101019344)

## 8. Table of contents

- This study presents a novel 4D bioprinting platform for engineering biomimetic musculoskeletal grafts.
- By tuning the mechanical properties of support baths, we enhance tissue fusion, collagen alignment, and cell differentiation.
- Using this strategy, we successfully fabricate scaled-up, anisotropic tissues such as meniscus, articular cartilage, and ligament
- This platform offers new solutions for advanced regenerative medicine applications.

