## Supplementary material for "A 4D Bioprinting Platform to Engineer Anisotropic Musculoskeletal Tissues by Spatially Patterning Microtissues into Temporally Adapting Support Baths": Supportin Information

### Supporting Information

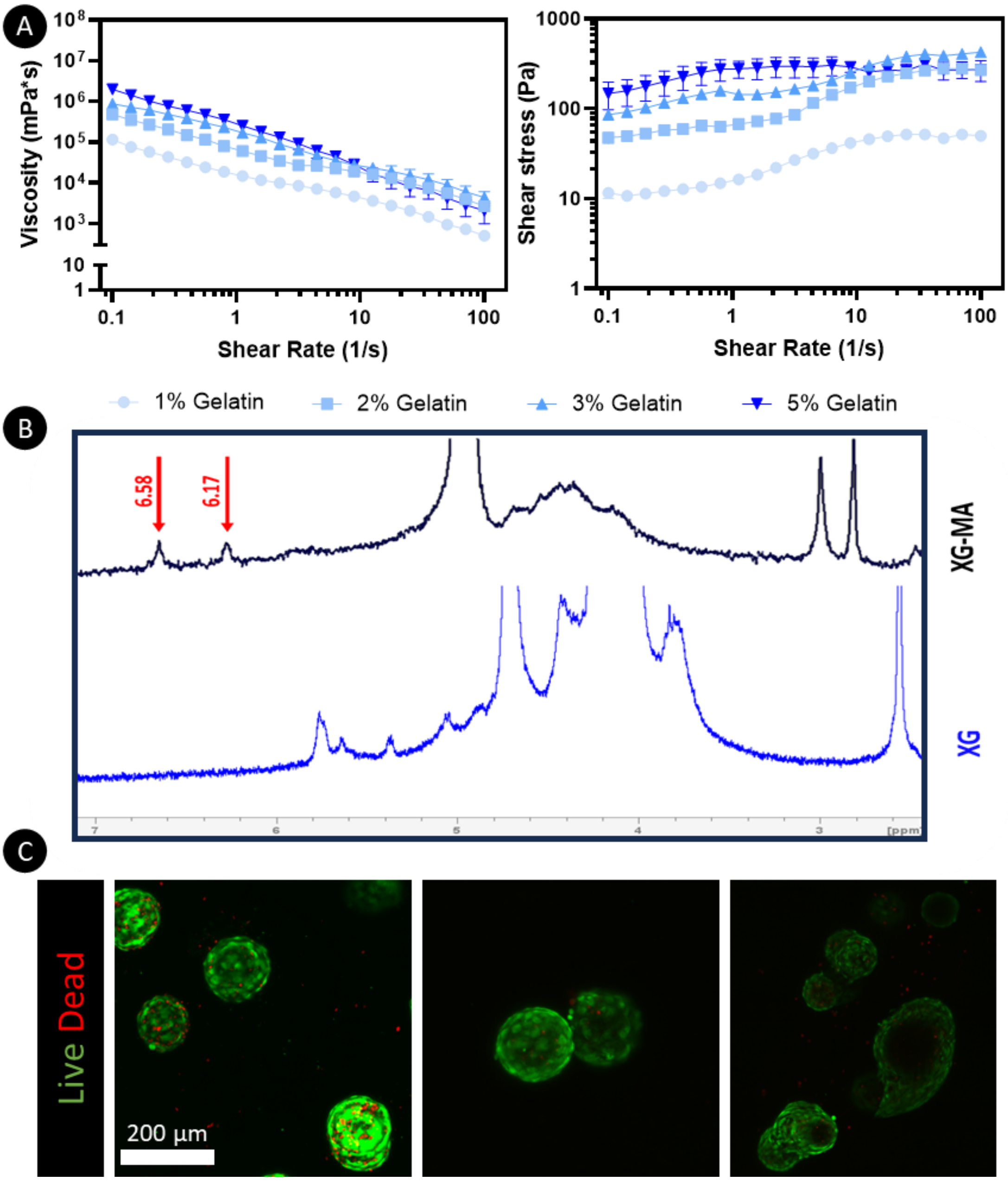


**Figure S1**. **Characterization of the XG-MA modification, gelatin viscoelastic behavior and cell viability, crucial for the successful development of the 4D bioprinting platform.** A) Viscoelastic behavior of gelatin bioink at different concentrations (5%, 3%, 2% and 1%) showing shear thinning behavior critical for bioink development. B) Proton nuclear magnetic resonance spectra ($^{1}$H-NMR) confirming the successful modification of XG with a methacrylic group (-C=C). Red arrows indicate the peaks of the methacrylic group at 6.58 and 6.17 ppm. C) Live/dead of microtissues (2000 cells/µT) encapsulated in gelatin 1% prior printing show high viability of microtissues. Green: Calcein (Live cells), Red: Ethidium Bromide (dead cells). Sb: 200 µm.

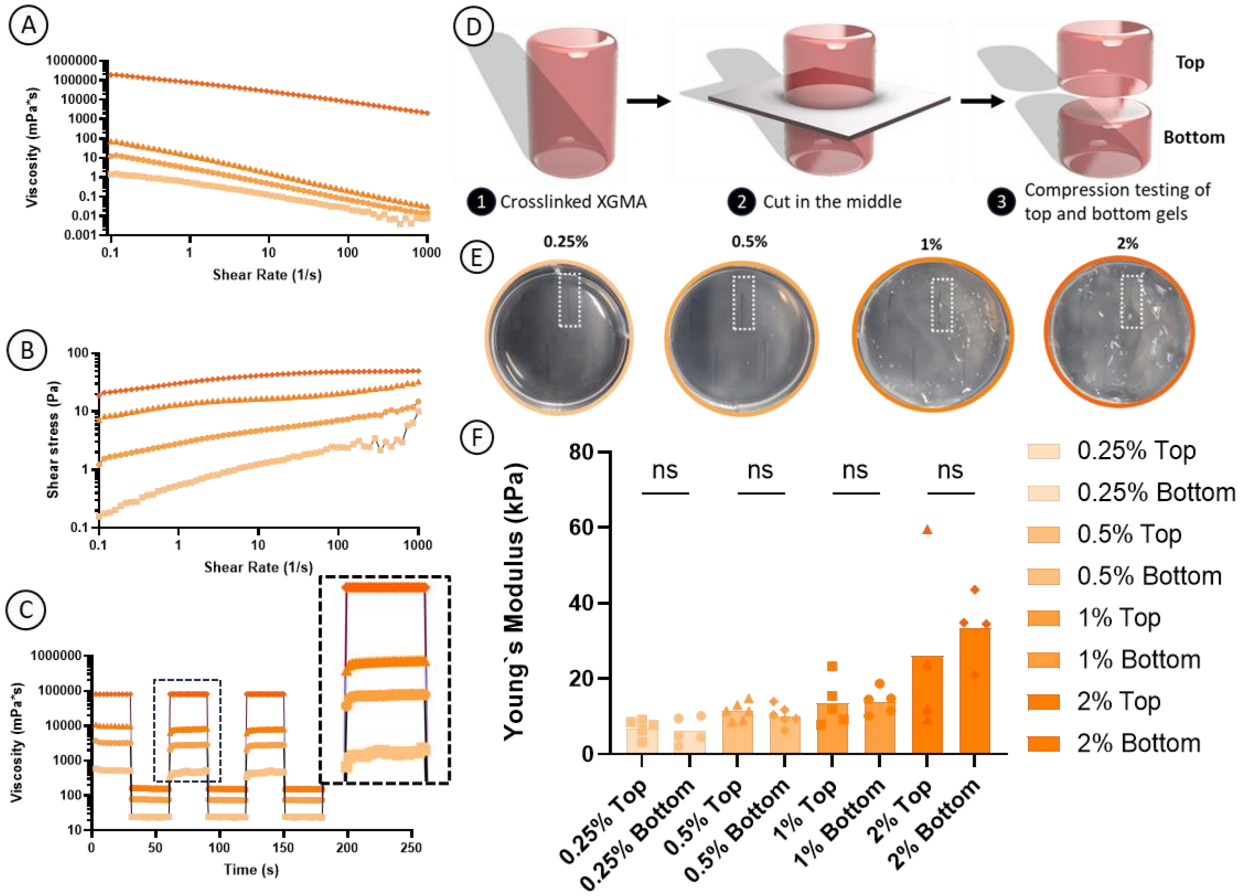


**Figure S2. Rheological properties of XG-MA at different concentrations and UV-curing depth influence on the bath stiffness.** A) Viscosity dependency on shear rate of the support bath at different concentrations, from low shear rate (0.1 $s^{-1}$) to very high shear stress (1000 $s^{-1}$) demonstrating shear-thinning properties B) Shear stress dependency on shear rate of the support bath at different concentrations, from low (0.1 $s^{-1}$) to high (1000 $s^{-1}$) shear stress C) Stress-relaxation behavior of the XG-MA support bath at different concentrations to assess the recovery of the material post applying high shear stress D) Graphical representation of mechanical testing of both halves (top and bottom) of the support bath post UV exposure to assess UV curing depth E) macroscopic pictures of acellular (1% gelatin only) printed filaments into support baths at different concentrations, with different optical properties F) mechanical testing of both halves (top and bottom) of each gel to assess curing depth dependency of all concentrations of the support bath post-UV exposure (4 min). A two-way ANOVA test was performed followed by Tukey's post comparison to compare the means of each group at different time points. Significance was accepted when $p<0.05$. ns: non-significant.

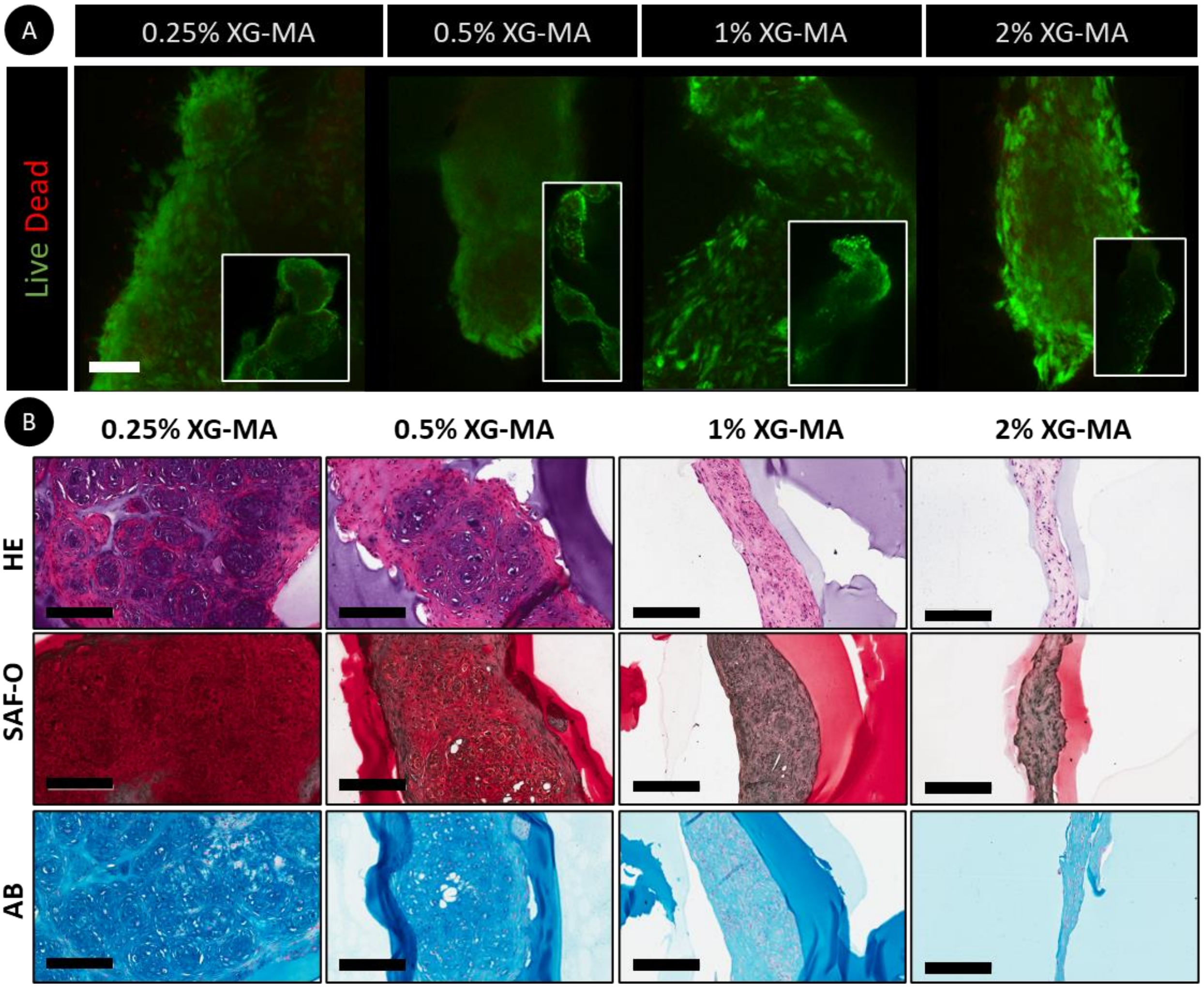


**Figure S3. Stiffness of the supporting bath post printing influences the deposition of sGAGs and matrix re(modelling) over time in culture.** A) Live/dead staining of printed microtissues into XG-MA support bath, 7 days after printing. Green: calcein AM; Red: Ethidium Bromide. Sb: 100µm. B) Histological evaluation of the printed microtissues after 28 days in culture with H&E, Safranin-O (SAF-O) and Alcian Blue to evaluate sGAGs deposition, which decreases with the increase in stiffness of the XG-MA support bath. Sb: 200 µm

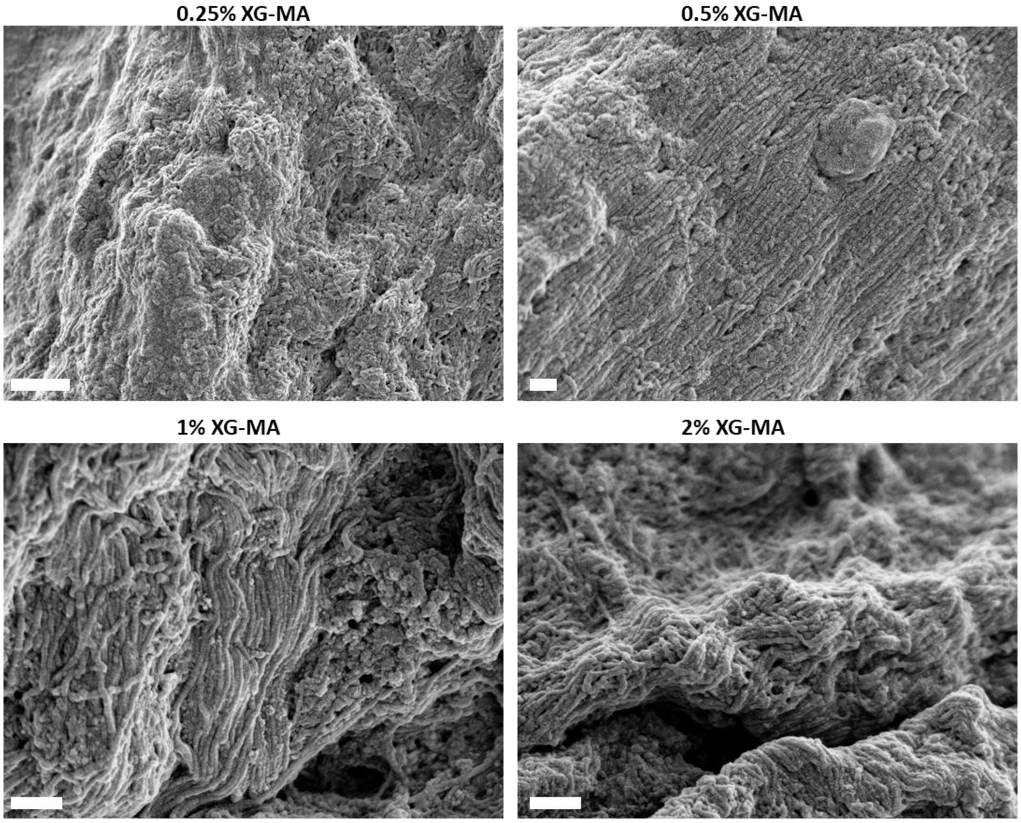


**Figure S 4. (Re)modelling of the collagen fibers along the printing direction after 28 days in culture.** Scanning electron microscope (SEM) pictures of bioprinted microtissues to show the ECM organization and (re)modelling, driving tissue maturation and differentiation. Sb: 10 µm.

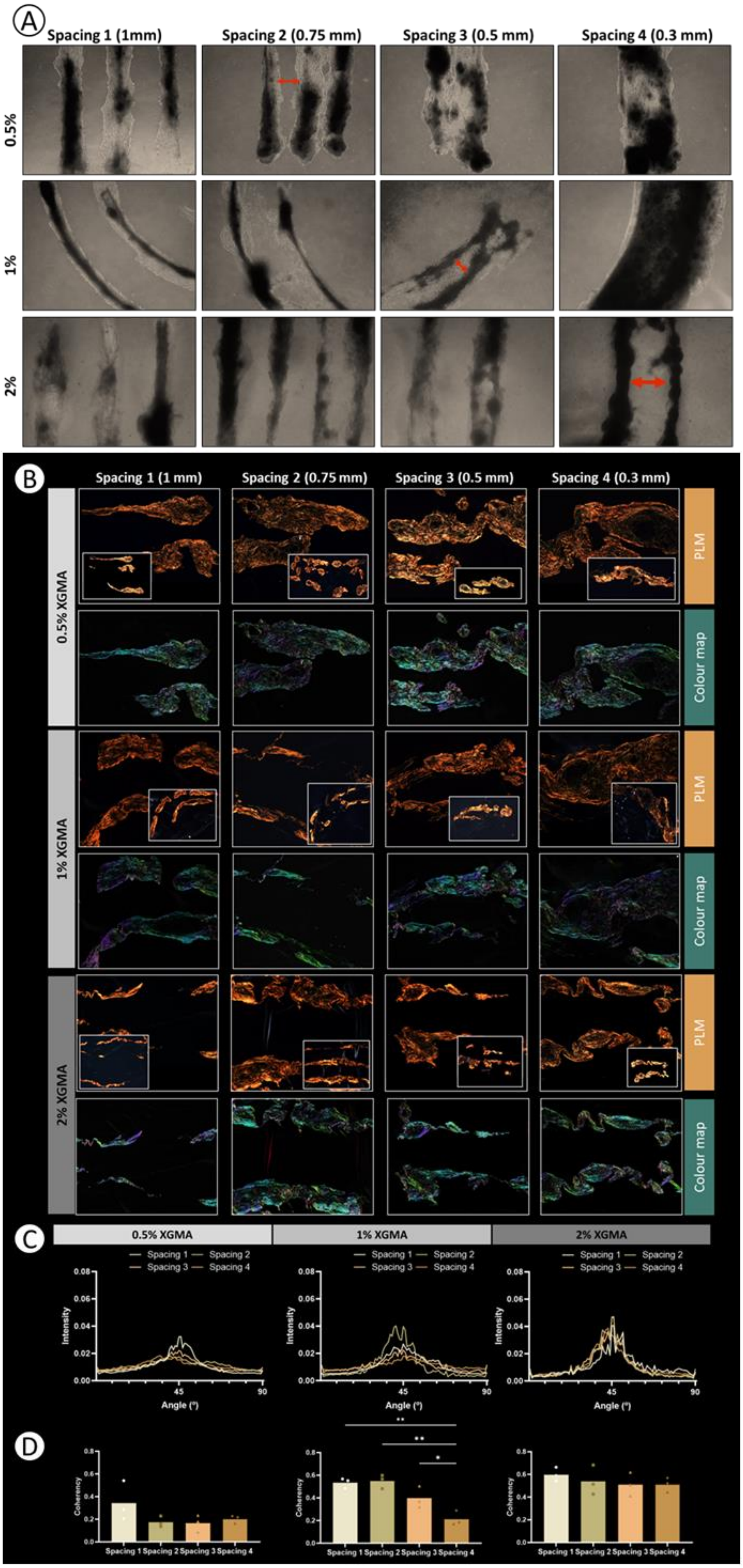
A
Spacing 1 (1mm)
Spacing 2 (0.75 mm)
Spacing 3 (0.5 mm)
Spacing 4 (0.3 mm)
0.5%
1%
2%
B
Spacing 1 (1 mm)
Spacing 2 (0.75 mm)
Spacing 3 (0.5 mm)
Spacing 4 (0.3 mm)
0.5% XGMA
1% XGMA
2% XGMA
PLM
Colour map
C
0.5% XGMA
1% XGMA
2% XGMA
Spacing 1
Spacing 2
Spacing 3
Spacing 4
Intensity
Angle (°)
D
Coherency
Spacing 1
Spacing 2
Spacing 3
Spacing 4

**Figure S5. The spacing (distance) between filaments post printing modulates the anisotropy of a scaled up bioprinted tissue construct**. A) Brightfield microscope pictures of bioprinted microtissues 28 days post printing showing fusion of the microtissues and adequate distance between each filament. B) Polarized Light Microscope images of bioprinted filaments at different distancing after 28 days of culture in TGF-β3 supplemented media, in support bath with variable stiffness. C) Mean average of the collagen fibers indicates that the spacing of the filaments into a scaled-up constructs influences the final anisotropy of the printed construct. D) Fiber coherency shows highly aligned tissues. Color maps are generated from PLM images. Here, color hue is used to indicate fiber orientation where blue/cyan indicated fibers oriented at 0 degrees and pink/red indicated fibers oriented at 90 degrees. For all the graphs a one-way ANOVA test was performed followed by Tukey's post comparison to compare the means of each group at different time points. Significance was accepted when $p<0.05$. * Indicates $p<0.05$. **Indicates $p<0.01$. For all the graphs: n=3.
